# Dengue Virus Antibodies Enhance Zika Virus Infection

**DOI:** 10.1101/050112

**Authors:** Lauren M. Paul, Eric R. Carlin, Meagan M. Jenkins, Amanda L. Tan, Carolyn M. Barcellona, Cindo O. Nicholson, Lydie Trautmann, Scott F. Michael, Sharon Isern

**Author notes:** Corresponding author (SI). Current Address: Department of Molecular Genetics and Microbiology, Duke University School of Medicine, Durham, North Carolina, United States of America (Cindo O. Nicholson). SI and SFM are Joint Senior Authors.

## Abstract

**Background:** For decades, human infections with Zika virus (ZIKV), a mosquito-transmitted flavivirus, were sporadic, associated with mild disease, and went underreported since symptoms were similar to other acute febrile diseases endemic in the same regions. Recent reports of severe disease associated with ZIKV, including Guillain-Barré syndrome and severe fetal abnormalities, have greatly heightened awareness. Given its recent history of rapid spread in immune naïve populations, it is anticipated that ZIKV will continue to spread in the Americas and globally in regions where competent *Aedes* mosquito vectors are found. Globally, dengue virus (DENV) is the most common mosquito-transmitted human flavivirus and is both well-established and the source of outbreaks in areas of recent ZIKV introduction. DENV and ZIKV are closely related, resulting in substantial antigenic overlap. Through a mechanism known as antibody-dependent enhancement (ADE), anti-DENV antibodies can enhance the infectivity of DENV for certain classes of immune cells, causing increased viral production that correlates with severe disease outcomes. Similarly, ZIKV has been shown to undergo ADE in response to antibodies generated by other flaviviruses. However, response to DENV antibodies has not yet been investigated.

**Methodology / Principal Findings:** We tested the neutralizing and enhancing potential of well-characterized broadly neutralizing human anti-DENV monoclonal antibodies (HMAbs) and human DENV immune sera against ZIKV using neutralization and ADE assays. We show that anti-DENV HMAbs, cross-react, do not neutralize, and greatly enhance ZIKV infection *in vitro*. DENV immune sera had varying degrees of neutralization against ZIKV and similarly enhanced ZIKV infection.

**Conclusions / Significance:** Our results suggest that pre-existing DENV immunity will enhance ZIKV infection *in vivo* and may increase disease severity. A clear understanding of the interplay between ZIKV and DENV will be critical in informing public health responses in regions where these viruses co-circulate and will be particularly valuable for ZIKV and DENV vaccine design and implementation strategies.

**Author Summary:** Recent reports of severe disease, including developmental problems in newborns, have greatly heightened public health awareness of Zika virus (ZIKV), a mosquito-transmitted virus for which there is no vaccine or treatment. It is anticipated that ZIKV will continue to spread in the Americas and globally in regions where competent mosquitoes are found. Dengue virus (DENV), a closely related mosquito-transmitted virus is well-established in regions of recent ZIKV introduction and spread. It is increasingly common that individuals living in these regions may have had a prior DENV infection or may be infected with DENV and ZIKV at the same time. However, very little is known about the impact of DENV infections on ZIKV disease severity. In this study, we tested the ability of antibodies against DENV to prevent or enhance ZIKV infection in cell culture-based assays. We found that DENV antibodies can greatly enhance ZIKV infection in cells.

Our results suggest that ZIKV infection in individuals that had a prior DENV infection may experience more severe clinical manifestations. The results of this study provide a better understanding of the interplay between ZIKV and DENV infections that can serve to inform public health responses and vaccine strategies.

## Introduction

Zika virus (ZIKV), a mosquito-transmitted flavivirus, was first isolated in a sentinel rhesus monkey and *Aedes africanus* mosquitoes in the Zika Forest near Entebbe, Uganda in 1947 during routine arbovirus surveillance by the Virus Research Institute in Entebbe [1]. A subsequent survey of human sera for ZIKV neutralizing antibodies in localities in Uganda including Zika, Kampala and Bwamba concluded that 6.1% of individuals tested were ZIKV seropositive [2]. Although no human disease had been associated with ZIKV at the time, it was speculated that ZIKV infection was not necessarily rare or unimportant. Neutralizing anti-ZIKV activity was found in serum collected between 1945 and 1948 from individuals residing in East Africa including Uganda and then northern Tanganyika south of Lake Victoria. Over 12% of individuals tested had ZIKV neutralizing activity though at the time ZIKV was an agent of unknown disease [3]. Simpson described the first well-documented case of ZIKV disease and virus isolation in humans [4]. He became infected while working in the Zika Forest in 1963, and his mild disease symptoms, that lasted for 5 days, included low-grade fever, headache, body aches, and a maculopapular rash. These symptoms have since become hallmark features of ZIKV human disease. In 1968, ZIKV was isolated from 3 non-hospitalized children in Ibadan, Nigeria indicating that ZIKV was not restricted to East Africa [5]. A 1953 and 1954 serological survey in South East Asia that included individuals from Malaysia near Kuala Lumpur, Thailand, and North Vietnam found ZIKV protective sera in individuals residing in these regions ranging from 75% positive in Malayans, 8% in Thailand, and 2% in North Vietnam [6]. An early 1980s serologic study of human volunteers in Lombok, Indonesia reported that 13% had neutralizing antibodies to ZIKV [7]. These studies illustrated that ZIKV had spread beyond Africa and at some point became endemic in Asia [8].

For decades, human ZIKV infections were sporadic, spread in geographic location, remained associated with mild disease, and perhaps went underreported since its symptoms were similar to other acute febrile diseases endemic in the same regions. As is the case with other flaviviruses, it is known that ZIKV antibodies cross-react with other flavivirus antigens including dengue virus (DENV) as was illustrated in the Yap State, Micronesia ZIKV outbreak in 2007. Initial serologic testing by IgM capture ELISA with DENV antigen was positive which led physicians to initially conclude that the causative agent for the outbreak was DENV, though the epidemic was characterized by a rash, conjunctivitis and arthralgia symptoms clinically distinct from DENV [9]. Subsequent testing using a ZIKV-specific reverse transcriptase polymerase chain reaction (RT-PCR) assay revealed that ZIKV was the causative agent [10]. Sequencing and phylogenetic analysis indicated that only one ZIKV strain circulated in the epidemic and that it had a 88.7% nucleotide and 96.5% amino acid identity to the African 1947 ZIKV strain MR766. A 12-nucleotide sequence was found in the envelope gene that was absent in the ZIKV African prototype. The consequence of this addition with regards to virus replication, fitness, and disease outcome is not yet known. No further transmission was reported in the Pacific until 2013 when French Polynesia reported an explosive ZIKV outbreak with 11% of the population seeking medical care [11]. Phylogenetic analysis revealed that the outbreak strain was most closely related to a Cambodia 2010 strain and the Yap State 2007 strain corroborating expansion of the Asian ZIKV lineage. Perinatal ZIKV transmission was also reported in French Polynesia [12]. In addition, 3% of blood bank samples tested positive for ZIKV by RT-PCR even though the donors were asymptomatic when they donated, underscoring the potential risk of ZIKV transmission through blood transfusions [13]. ZIKV transmission and spread maintained a solid foothold in the Pacific [14] and continued its spread in 2014 with confirmed outbreaks in French Polynesia, New Caledonia, Easter Island, and the Cook Islands [15-18].

The first report of local transmission of ZIKV in the Americas occurred in the city of Natal in Northern Brazil in 2015 [19]. Natal patients reported intense pain resembling Chikungunya virus (CHIKV) infection but with a shorter clinical course, in addition to maculopapular rash. No deaths or complications were reported at the time, though given the naïve immunological status of the Brazilian population, ZIKV expansion was predicted. Several theories arose to explain the probable introduction of ZIKV into Brazil. These included the soccer World Cup in 2014, though no ZIKV endemic countries competed [19], the 2014 Va’a World Sprint Championships canoe race held in Rio de Janeiro with participants from French Polynesia, New Caledonia, Cook Islands, and Easter Island [20], and the 2013 Confederations Cup soccer tournament which included competitors from French Polynesia [21]. Molecular clock analysis of various Brazilian ZIKV strains estimated that the most recent common ancestor dated back to 2013 making the first two theories less likely [21]. By mid-January 2016, ZIKV transmission had occurred in 20 countries or territories in the Americas as reported to the Pan American Health Organization [22]. The primary mode of ZIKV transmission appeared to be through mosquito vectors, although cases of perinatal and sexual transmission were also reported [12,23]. Given its recent history of rapid spread in immune naïve populations, it is anticipated that ZIKV will continue to spread for the foreseeable future in the Americas and globally in regions where competent *Aedes* mosquito vectors are present. Kindhauser et al. 2016 can serve as a comprehensive account of the world-wide temporal and geographic distribution of ZIKV from 1947 to present day [24].

Until relatively recently, due to its mild clinical outcome, ZIKV disease had not been a critical public health problem. As a result, compared to other related viruses, it remained understudied. However, recent reports of severe ZIKV disease including Guillain-Barré syndrome in French Polynesia [14,25] and associations between ZIKV and microcephaly and other severe fetal abnormalities in Brazil [26-30] have greatly heightened awareness of ZIKV. Retrospectively, the incidence of Guillain-Barré syndrome during the 2014 ZIKV French Polynesia outbreak and the incidence of microcephaly in Brazil in 2015 were both 20 times higher than in previous years. The cause of these severe ZIKV disease outcomes remains an open question. Recent ZIKV outbreaks in the Pacific and the Americas have been explosive and associated with severe disease, yet earlier expansions in Africa and Asia were gradual, continuous and associated with mild clinical outcomes. Much of the difference may lie in the age of exposure. In ZIKV endemic areas, most adults have pre-existing ZIKV immunity and new cases primarily occur in children. Introduction of ZIKV into immune naïve populations where all ages are susceptible to infection, including women of child-bearing age, is the new scenario for ZIKV expansion. However, we are still left without an understanding of why certain individuals develop severe disease such as Guillain-Barré syndrome, and why some expectant mothers transmit ZIKV to their developing offspring *in utero*, resulting in fetal infection and developmental abnormalities, whereas others do not. A possible explanation could be the impact of pre-existing immunity to co-circulating flaviviruses.

Globally, DENV is the most common mosquito-transmitted human flavivirus [31] and is both well-established and the source of new outbreaks in many areas of recent ZIKV introduction [15,16]. DENV and ZIKV are very closely related resulting in substantial antigenic overlap. The four serotypes of DENV (DENV-1, DENV-2, DENV-3, and DENV-4) have an antigenic relationship that impacts disease severity. Infection with one serotype typically results in a life-long neutralizing antibody response to that serotype, but yields cross-reactive, non-neutralizing antibodies against the other serotypes. These cross-reactive, non-neutralizing antibodies are responsible for antibody-dependent enhancement (ADE), a phenomenon where DENV particles are bound (opsonized) by these antibodies, which allows the infection of antibody Fc receptor (FcR) bearing cells, such as macrophages, dendrocytes, and monocytes, that are normally not infected. The presence of enhancing antibodies correlates with increased DENV viremia and disease severity [32-34]. Similarly, ZIKV has also been shown to undergo ADE in response to sub-neutralizing concentrations of homologous anti-serum, and in response to heterologous anti-serum from several different flaviviruses [35]. In addition, anti-ZIKV sera has been shown to enhance infectivity of related viruses [36]. In one study, immune mouse ascites against various flaviviruses including ZIKV, West Nile virus (WNV), Yellow Fever-17D (YF17D), Wesselsbron virus, Potiskum, Dakar Bat, and Uganda S were tested for ZIKV ADE in P388D_1_, a mouse macrophage Fc receptor cell line [35]. All heterologous immune mouse ascites, as well as homologous immune ascites, enhanced ZIKV in culture. Of note, the fold-enhancement was greater for ZIKV compared to peak enhancement of other flaviviruses tested against their heterologous immune ascites. Given the incidence of co-circulating flaviviruses, the study authors alluded to the importance of testing human sera for ADE potential of circulating flaviviruses. In a subsequent study, human cord blood and sera of newborns and adults collected in Ibadan, Nigeria, was tested for ADE of DENV-2, YF17D and WNV in P388D_1_, but the ADE potential of ZIKV was not tested [37]. To our knowledge, only mouse sera and mouse cells have been used to date for *in vitro* ZIKV ADE assays. In addition, anti-DENV immune serum has never been tested for ZIKV enhancement activity. Curiously, the 2013-14 French Polynesia ZIKV outbreak demonstrated that all the patients with Guillain-Barré syndrome had pre-existing DENV immunity [25].

In this study, we investigated the role that pre-existing DENV immunity plays during ZIKV infection. Here we report that human anti-DENV serum and well-characterized human anti-DENV monoclonal antibodies (HMAbs) cause substantial ZIKV ADE in a human Fc receptor bearing cell line. Our results suggest that pre-existing antibodies from a prior DENV infection will enhance ZIKV infection *in vivo* and may increase disease severity.

## Methods

### Human Sera and Monoclonal Antibodies

The collection of human blood samples was reviewed and approved by the institutional review board of Florida Gulf Coast University (protocols 2007-08 and 2007-12) and the research ethics committee of the Centre Hospitalier de l’Université de Montréal. Informed written consent was obtained from all subjects. Jamaica 1, and Singapore 1 sera have been previously described, from subject 8C and subject DA003, respectively [38]. Subject Jamaica 1 (8C) was infected with DENV in Jamaica in 2007 and had blood drawn in 2008, approximately 3 months post-recovery. The subject had fever for 12 days, headache, retro-orbital pain, and blood in sputum. Subject Jamaica 2 (10E) was infected with DENV in Jamaica in 2007 with severe symptoms and had blood drawn in 2008, 3 months after recovery. Subject Singapore 1 (DA003) was hospitalized in Singapore in 2008 for complications due to DENV infection and had blood drawn approximately 4 weeks post-recovery. No hemoconcentration or bleeding was present. Subject Singapore 2 (PHC) was infected with DENV and hospitalized in Singapore in 2008 and had blood drawn approximately 4 weeks after recovery. A healthy subject from Montreal, Canada provided control serum that was collected in 2003 prior to vaccination with yellow fever 17D vaccine. Travel history confirmed that the subject had not travelled to regions outside North America and had no previous exposure to DENV or ZIKV. Sera were heat inactivated for 30 min at 56°C prior to use. Anti-DENV HMAbs 1.6D and D11C isolated from subject Jamaica 1 and Singapore 1, respectively, were kindly provided by J. S. Schieffelin from Tulane University and have been well-characterized and described previously [38].

### Viruses and Cell Culture

The 1947 Ugandan isolate, ZIKV MR766, and DENV-1 strain HI-1, DENV-2 strain NG-2, DENV-3 strain H-78, and DENV-4 strain H-42, were kindly provided by R. B. Tesh at the University of Texas at Galveston through the World Reference Center for Emerging Viruses and Arboviruses. ZIKV stock was propagated by single passage in African green monkey (*Cercopithecus aethiops*) kidney epithelial cells, Vero (ATCC CCL-81, American Type Culture Collection, Manassas, VA), cultured in Eagle’s Minimum Essential Medium supplemented with 10% (v/v) fetal bovine serum (FBS), 2mM Glutamax, 100U/mL penicillin G, 100ug/mL streptomycin, and 0.25ug/mL amphotericin B at 37°C with 5% (v/v) CO_2_. Rhesus macaque (*Macaca mulatta*) kidney epithelial cells, LLC-MK2 (ATCC CCL-7) used to propagate DENV and titer and perform focus-forming unit reduction neutralization assays, were cultured in Dulbecco’s Modified Eagle Medium (DMEM) supplemented with 10% (v/v) FBS, 2mM Glutamax, 100U/mL penicillin G, 100ug/mL streptomycin, and 0.25ug/mL amphotericin B at 37°C with 5% (v/v) CO_2_. Human bone-marrow lymphoblast cells bearing FcRII, K-562 (ATCC CCL-243) used to perform antibody-dependent enhancement assays (ADE), were cultured in RPMI-1640 (Hyclone, Logan, UT) supplemented with 10% (v/v) FBS, 2mM Glutamax, 100U/mL penicillin G, 100ug/mL streptomycin, and 0.25ug/mL amphotericin B at 37°C with 5% (v/v) CO_2_. All reagents were purchased from ThermoFisher, Waltham, MA unless otherwise noted.

### Enzyme-linked Immunosorbent Assay

Enzyme-linked immunosorbent assays (ELISA) were performed as follows. Corning brand high-bind 96-well plates (ThermoFisher, Waltham, MA) were coated with 100uL Concanavalin A (ConA) (Vector Laboratories, Burlingame, CA) at 25ug/mL in 0.01M HEPES (Sigma, Saint Louis, MO) and incubated for 1 hr at room temperature. Wells were washed with phosphate buffered saline (PBS) with 0.1% (v/v) Tween 20 (Sigma) and incubated for 1 hr at room temperature with 100uL of filtered ZIKV or DENV-2 culture supernatant inactivated with 0.1% (v/v) Triton-X100 (Sigma). After a wash step with PBS containing 0.1% (v/v) Tween 20, wells were blocked with 200uL PBS containing 0.5% (v/v) Tween 20 and 5% (w/v) non-fat dry milk for 30 min. Primary HMAbs D11C and 1.6D in PBS containing 0.5% (v/v) Tween 20 were incubated for 30 min at room temperature. After a wash step, 100uL of a peroxidase-conjugated affinity purified anti-human IgG (Pierce, Rockford, IL) diluted to 1ug/mL in PBS-0.5% (v/v) Tween 20 was incubated at room temperature for 30 min to detect the primary antibody. After a final wash step, color was developed with tetramethylbenzidineperoxide (ProMega, Madison, WI) as the substrate for peroxidase. The reaction was stopped after 3 min by adding 100uL1M phosphoric acid (Sigma), and the absorbance was read at 450 nm. Negative controls included media without virus, ConA only, and ConA without primary or secondary antibodies.

### Focus-forming Assay

Focus-forming assays were performed essentially as previously described [38]. LLC-MK2 target cells were seeded at a density of approximately 500,000 cells in each well of a 12-well plate 24-48 hrs prior to infection. For titer assays, 10-fold serial dilutions of virus were prepared. For neutralization assays, approximately 100 focus-forming units of virus were incubated with dilutions of heat-inactivated serum or purified HMAbs in serum-free DMEM for 1 hr at 37°C. Mixtures were allowed to infect confluent target cell monolayers for 1 hr at 37°C, with rocking every 15 min, after which the inoculum was aspirated and cells were overlaid with fresh Minimum Essential Medium (MEM) supplemented with 10% (v/v) FBS, 2mM Glutamax, 100U/mL penicillin G, 100ug/mL streptomycin, and 0.25ug/mL amphotericin B containing 1.2% (w/v) microcrystalline cellulose Avicel (FMC BioPolymer, Philadelphia, PA). The infected cells were then incubated at 37°C with 5% (v/v) CO_2_ for 48 hr (DENV-4), 60 hr (ZIKV), or 72 hr (DENV-1, −2, and −3). Cells were fixed in Formalde-Fresh Solution (ThermoFisher), either overnight at 4°C or for 1 hr at room temperature and permeabilized with 70% (v/v) ethanol for 30 min. Foci were detected using primary HMAbs 1.6D or D11C incubated for 8 hr at room temperature, followed by secondary horseradish peroxidase-conjugated goat anti-human IgG (H+L) (Pierce, Rockford, IL) incubated for 8 hr at room temperature. Foci were visualized by the addition of 3,3-diaminobenzidine tetrahydrochloride (Sigma-Aldrich, St. Louis, MO).

### Antibody-dependent Enhancement Assay

Antibody-dependent enhancement assays were performed as previously described [38,39]. Briefly, 250 focus-forming units of ZIKV were mixed with human sera or HMAbs and RPMI medium in a 200ul volume and incubated for 1 hr at 37°C. Mixtures were added to 80,000 K562 cells in 300ul of complete RPMI medium and incubated for 3 days at 37°C, 5% (v/v) CO_2_. Control experiments were performed by pre-incubating cells for 1 hr at 37°C with a mouse anti-human FcRII MAb (anti-CD32) (Biolegend, San Diego, CA). Cells were collected by centrifugation and total RNA was isolated using an RNeasy Mini-kit (Qiagen, Valencia, CA) following the manufacturer’s protocol. Quantitative reverse transcription (qRT-PCR) was performed on isolated RNA using ZIKV-specific forward (CTGCTGGCTGGGACACCCGC) and reverse (CGGCCAACGCCAGAGTTCTGTGC) primers to amplify a 99bp product from the ZIKV NS5 region. A Roche LightCycler 480 II was used to run qRT-PCR using a LightCycler RNA Master SYBR Green I kit (Roche, Indianapolis, IN). Amplification conditions were as follows: reverse transcription at 61°C for 40 min, denaturation at 95°C for 30 sec, followed by 45 cycles of denaturing at 95°C for 5 sec, annealing at 47°C for 10 sec, and extension at 72°C for 15 sec.

## Results

### Cross-recognition of ZIKV E protein by human anti-DENV antibodies

It is well known that infection with closely related flaviviruses often results in a cross-reactive serum antibody response. The primary neutralizing epitopes targeted by human antibodies during a flavivirus infection are found in the envelope glycoprotein (E protein) [38,40-46]. The role of the E protein is to facilitate virus entry by binding and mediating the fusion of the virus membrane and cellular membrane in target cells. The E protein of ZIKV and the four serotypes of DENV have a high degree of genetic similarity and the amino acid sequence of fusion loop region of these viruses is identical. In a previous study, we characterized broadly neutralizing anti-DENV human monoclonal antibodies (HMAbs) derived from patients that had recovered from DENV infection [38]. These HMAbs recognized the E protein with high affinity, neutralized the four DENV serotypes, and mediated ADE *in vitro* at subneutralizing concentrations. Their neutralization activities correlated with a strong inhibition of intracellular fusion, rather than virus-cell binding. Additionally, we mapped epitopes of these HMAbs to the highly conserved fusion loop region of the E protein.

Given the high degree of similarity between the DENV E protein and the ZIKV E protein, we thus tested the ability of two of these well-characterized anti-DENV HMAbs, 1.6D and D11C, to recognize the glycosylated ZIKV E surface protein using a conA capture assay [38]. In this assay, the glycoprotein-binding lectin, conA, is used to capture ZIKV MR766 E glycoprotein, which is then recognized by anti-DENV HMAbs that recognize the DENV E protein fusion loop. The HMAb is then detected with an anti-human IgG HRP-conjugated secondary antibody and an HRP colorimetric substrate. Our results show that anti-DENV HMAbs, 1.6D and D11C, strongly recognize the ZIKV E surface glycoprotein (**Fig 1A, B**). In addition, we tested the ability of these HMAbs to recognize ZIKV-infected cells in an immunostained focus forming assay (**Fig 1C, D**). This result confirms that anti-DENV E fusion loop HMAbs cross-react with ZIKV.

**Fig 1.**
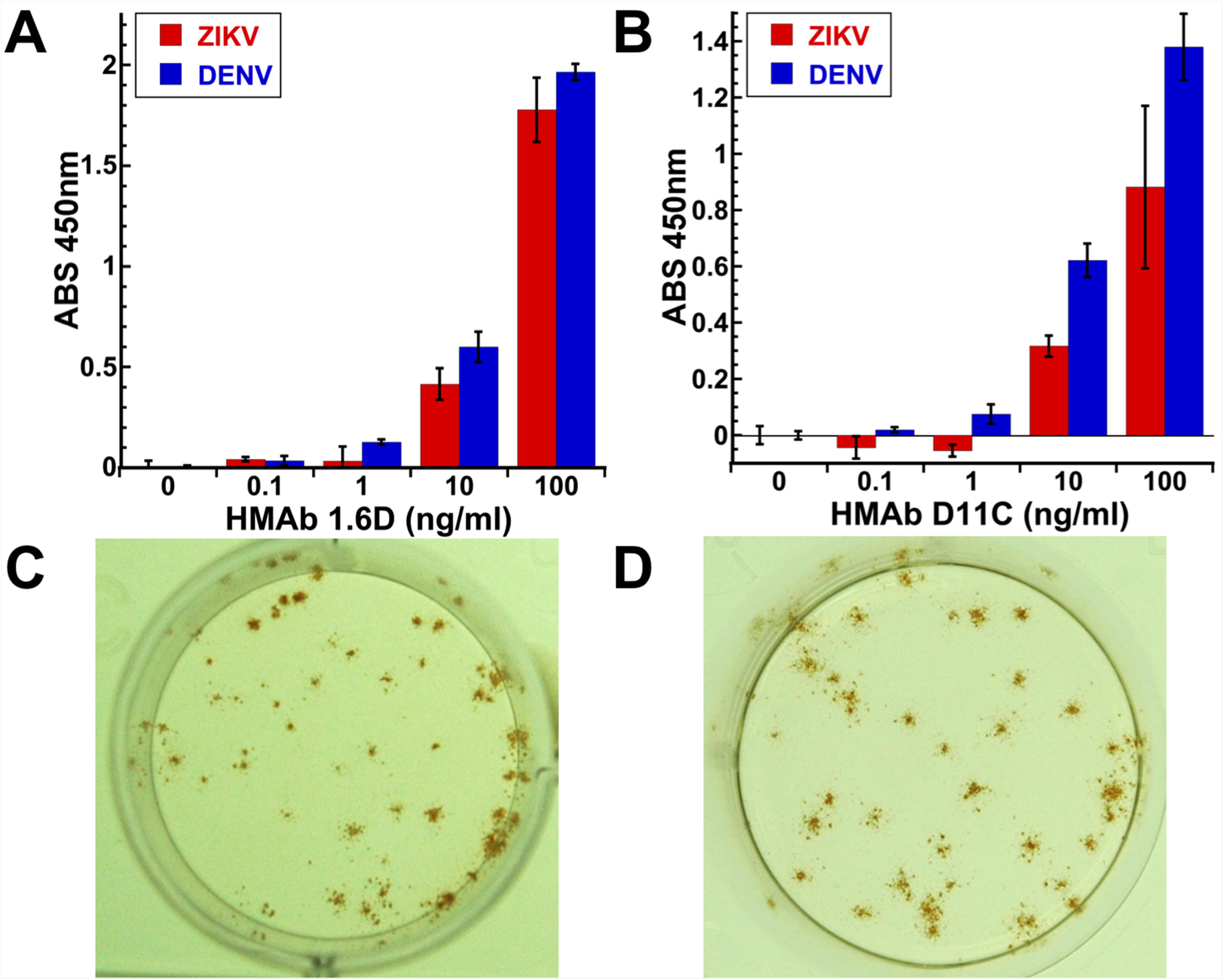
Cross-reactivity of anti-DENV HMAbs against ZIKV. Anti-DENV HMAbs 1.6D and D11C that recognize the DENV E protein fusion loop cross-react with ZIKV MR766 strain E surface glycoprotein as shown by ELISA (**A** 1.6D, **B** D11C) and recognize ZIKV infected cells in an immunostained focus-forming assay (**C** 1.6D, **D** D11C). DENV E is serotype 2, strain NG-2. Data shown are representative of two independent assays each done in triplicate.

### *In vitro* ZIKV neutralization activity of broadly neutralizing anti-DENV HMAbs

Since anti-DENV HMAbs 1.6D and D11C were cross-reactive against ZIKV, we tested whether they could neutralize ZIKV infectivity using an immunostained focus-forming unit reduction neutralization assay in rhesus macaque LLC-MK2 kidney epithelial cells [38]. Fusion loop HMAbs D11C and 1.6D are broadly neutralizing against all four DENV serotypes and represent a very common class of broadly neutralizing HMAbs, perhaps the dominant broadly neutralizing class of antibodies against DENV [38]. However, neither 1.6D nor D11C inhibited ZIKV infectivity *in vitro* at the concentrations tested (up to 40 ug/ml) (**Fig 2**). Broadly neutralizing anti-DENV HMAbs that target the E protein fusion loop bind to ZIKV antigens, but do not neutralize infectivity.

**Fig 2.**
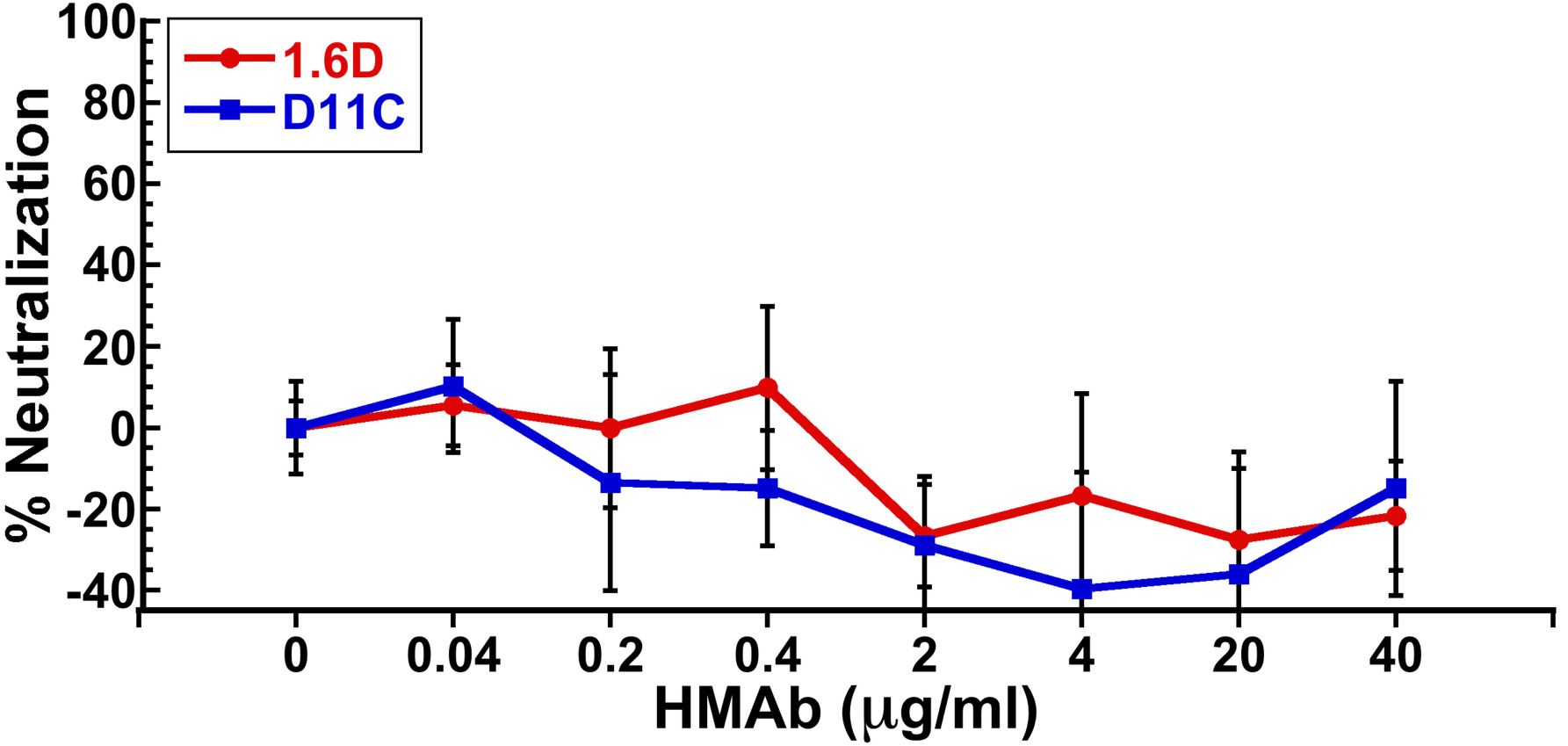
Neutralizing activity of anti-DENV HMAbs against ZIKV. Broadly neutralizing anti-DENV HMAbs 1.6D and D11C do not inhibit ZIKV MR766 infection in LLC-MK2 cells at the concentrations tested. The results shown are the average +/- the standard deviation of 6 replicates.

### *In vitro* ZIKV enhancement activity of broadly neutralizing anti-DENV HMAbs

DENV antibody-dependent enhancement (ADE) of Fc receptor (FcR)-bearing cells, which include macrophages, monocytes, and dendrocytes, correlates with increased viremia and severe disease outcomes [47]. Antibodies that recognize DENV surface proteins, but do not neutralize infectivity, can direct viral binding and infection of certain FcR cells that are not normally infected. Since anti-DENV HMAbs 1.6D and D11C cross-reacted with ZIKV proteins, but did not neutralize ZIKV infection, we tested whether they could mediate ZIKV ADE *in vitro*. In **Fig 3**, we show that ZIKV infection of FcR-bearing K562 cells can be strongly enhanced by anti-DENV HMAbs 1.6D (~140-fold) and D11C (~275-fold).

**Fig 3.**
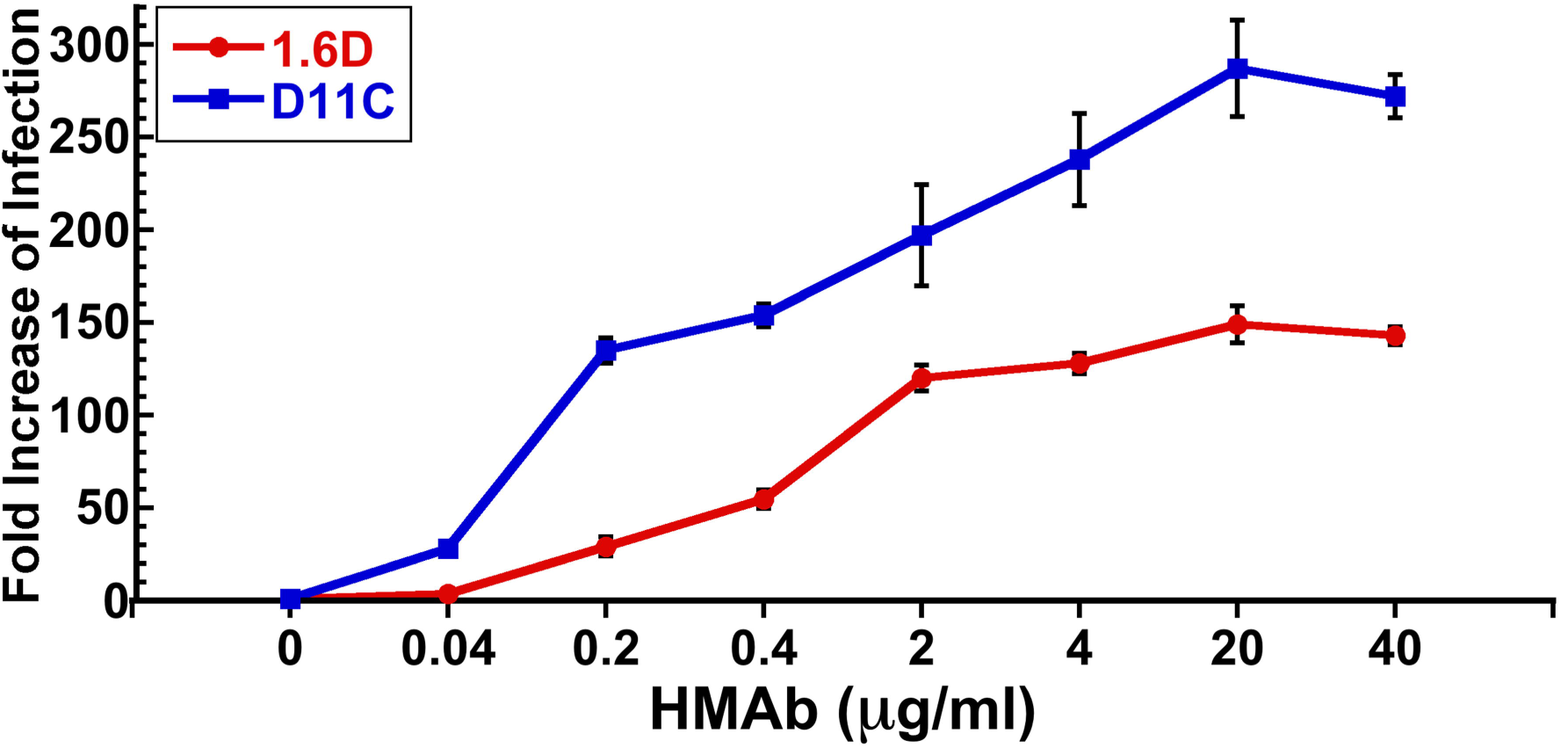
Enhancing activity of anti-DENV HMAbs against ZIKV. Broadly neutralizing anti-DENV HMAbs 1.6D and D11C show strong ZIKV MR766 infection enhancing activity. Independent assays were repeated twice in triplicate.

### *In vitro* ZIKV neutralization activity of human anti-DENV serum

Given the cross-reactive and strongly enhancing potential of anti-DENV HMAbs 1.6D and D11C, we investigated whether immune sera from DENV recovered patients contained other types of antibodies that could neutralize ZIKV infection. For this study, we wanted to investigate what might be considered the ‘worst case scenario’ with regards to pre-existing immunity to DENV. We selected sera from individuals with probable secondary DENV infection that had been collected in countries where multiple serotypes of DENV have been known to circulate. This scenario would serve to model the immune status of many individuals in regions where ZIKV is rapidly spreading.

We tested two human anti-DENV sera from Singapore and two from Jamaica, in addition to serum from a DENV-negative donor from Canada. The Singapore patient sera were collected in 2008 during which time ZIKV was endemic in Southeast Asia and after its expansion in the Yap State in Micronesia in the Pacific in 2007. The Canada donor serum was collected in 2003 and the Jamaica sera were collected in 2008 prior to documented introduction of ZIKV in the Americas. Additionally, the Jamaica and Canada subjects had no travel history to ZIKV endemic countries. We purposely selected Singapore 1 and Jamaica 1 sera for these studies since subject Singapore 1 was the source of HMAb D11C and subject Jamaica 1 was the source of HMAb 1.6D [38]. We wanted to determine whether the antibody repertoire of the same individuals contained DENV antibodies that could also neutralize ZIKV infection. Singapore 2 and Jamaica 2 sera were selected based on their broadly neutralizing activity against DENV. As shown in **Fig 4**, the Singapore (1 and 2) and Jamaica (1 and 2) sera showed broadly neutralizing activity against all four serotypes of DENV [38], indicating that they were likely from subjects with secondary DENV infections.

**Fig 4.**
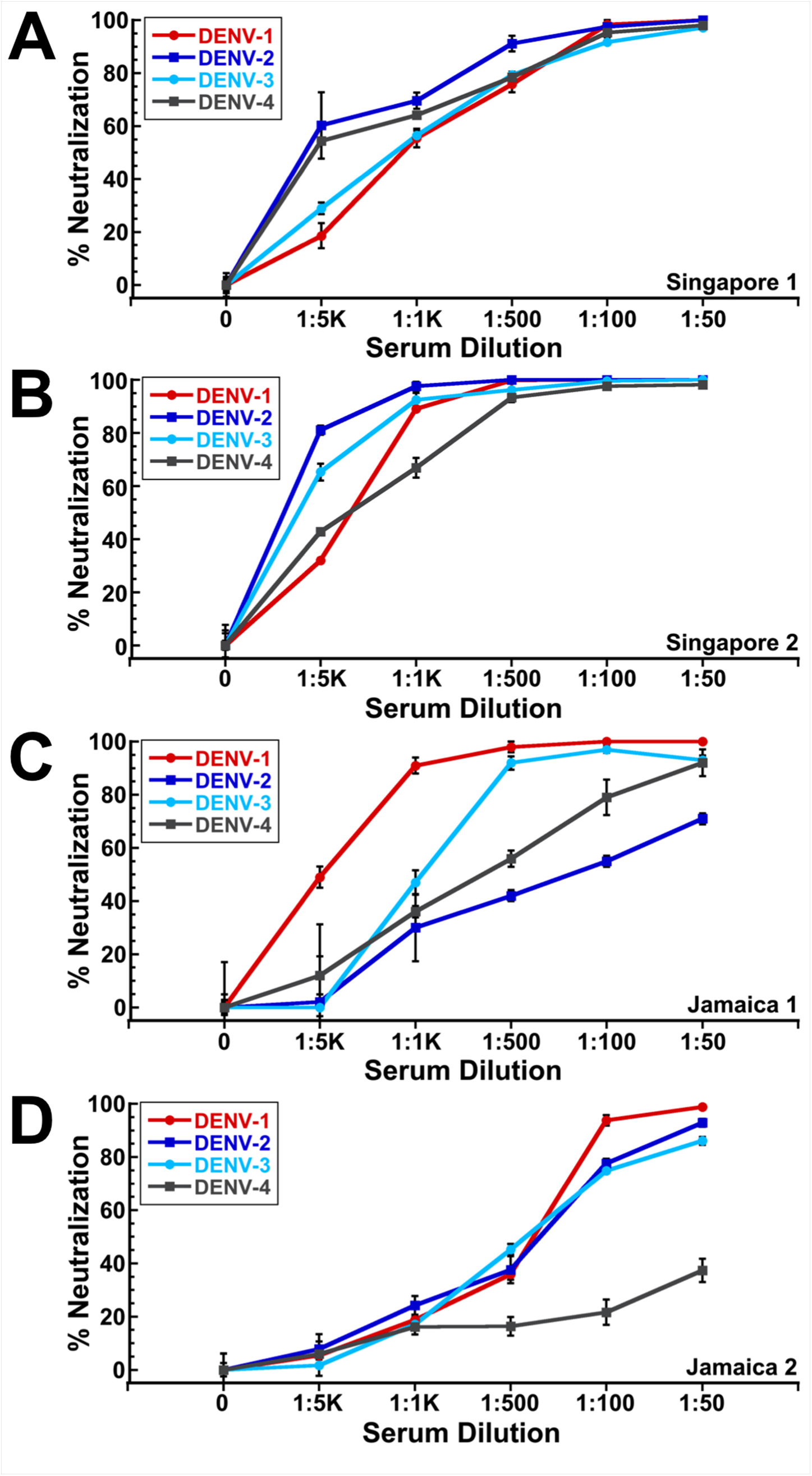
Neutralizing activity of anti-DENV human sera against DENV. All anti-DENV human sera showed broad neutralizing activity against multiple DENV serotypes 1-4. (**A**) Singapore 1, (**B**) Singapore 2, (**C**) Jamaica 1, (**D**) Jamaica 2. DENV-1, −2, −3, and −4 neutralizing activity of Singapore 1 and Jamaica 1 sera has previously been described and is shown here for clarity [38].

The results of the ZIKV neutralization assays with human anti-DENV sera are shown in **Fig 5**. We found that Singapore 1 serum strongly neutralized ZIKV, even at high dilutions (1:10,000 dilution), while Singapore 2 had no ZIKV neutralizing activity. Jamaica 1 serum neutralized ZIKV at the highest serum concentrations tested (1:100, 1:50), while Jamaica 2 serum did not. We suspect that the strongly ZIKV neutralizing Singapore 1 serum may be the result of a prior undiagnosed ZIKV infection, as ZIKV has been present in Southeast Asia for decades [6,7,24]. However, the less potent neutralizing activity from Jamaica 1 serum is very likely due to cross-neutralization from prior DENV infection, or infections, as ZIKV was unknown in the Americas at the time the serum was collected. Serum from Canada with no exposure to DENV or ZIKV was used as a negative control and had no ZIKV neutralizing activity [48].

**Fig 5.**
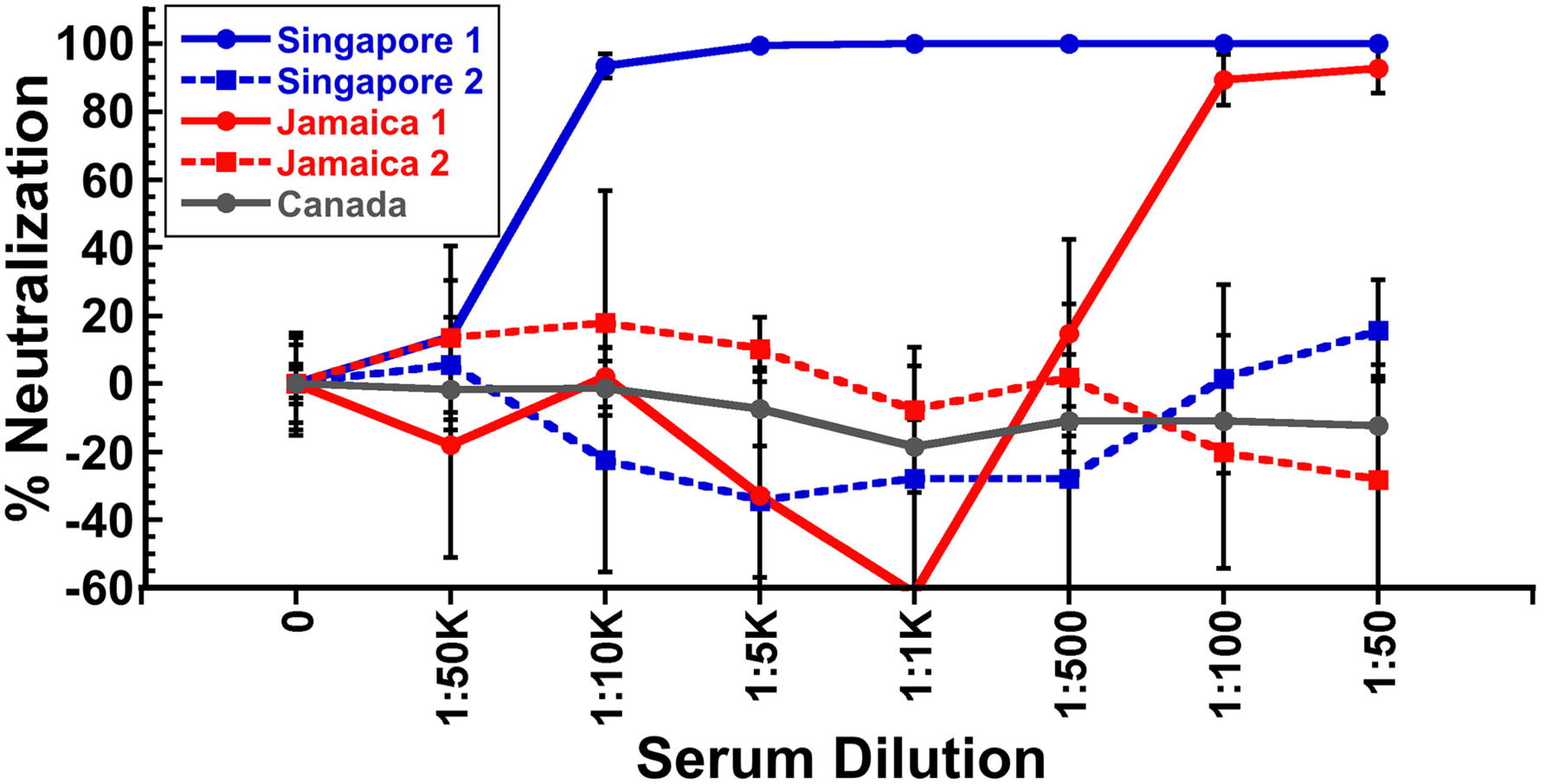
Neutralizing activity of anti-DENV human sera against ZIKV. Human anti-DENV sera from Singapore and Jamaica show both non-neutralizing and neutralizing activity against ZIKV MR766. Singapore 1 serum strongly neutralizes ZIKV MR766, suggesting prior ZIKV infection, while Singapore 2 serum has no neutralizing activity. Jamaica 1 serum neutralizes ZIKV MR766 at high serum concentrations, while Jamaica 2 serum shows no neutralizing activity at the dilutions tested. Control serum from Canada shows no ZIKV neutralizing activity. The results shown are the average +/- the standard deviation of 6 replicates.

### *In vitro* ZIKV enhancement activity of human anti-DENV serum

We then tested whether human DENV immune sera could mediate ADE *in vitro*. We show that ZIKV infection of FcRII bearing K562 cells can be strongly enhanced (up to 200 fold) by all human anti-DENV sera tested (**Fig 6**). In comparison, the control serum from Canada showed no enhancement. The highly neutralizing Singapore 1 serum showed strong ZIKV enhancement at intermediate dilutions (1:100,000 to 1:10,000) that diminished at lower dilutions (1:5,000 to 1:100), indicating that highly neutralizing antibodies can overcome ZIKV infection enhancement at sufficiently high concentrations. To confirm that the mechanism of enhancement involved entry of antibody-bound ZIKV particles through the K562 FcRII pathway, we pre-incubated K562 cells with a mouse anti-FcRII MAb prior to infection with ZIKV that had been pre-incubated with a highly enhancing dilution (1:50,000) of the ZIKV-neutralizing Singapore 1 serum. Our results demonstrate that the ZIKV enhancement effect can be effectively blocked in a dose-responsive manner with an anti-FcRII MAb (**Fig 7**).

**Fig 6.**
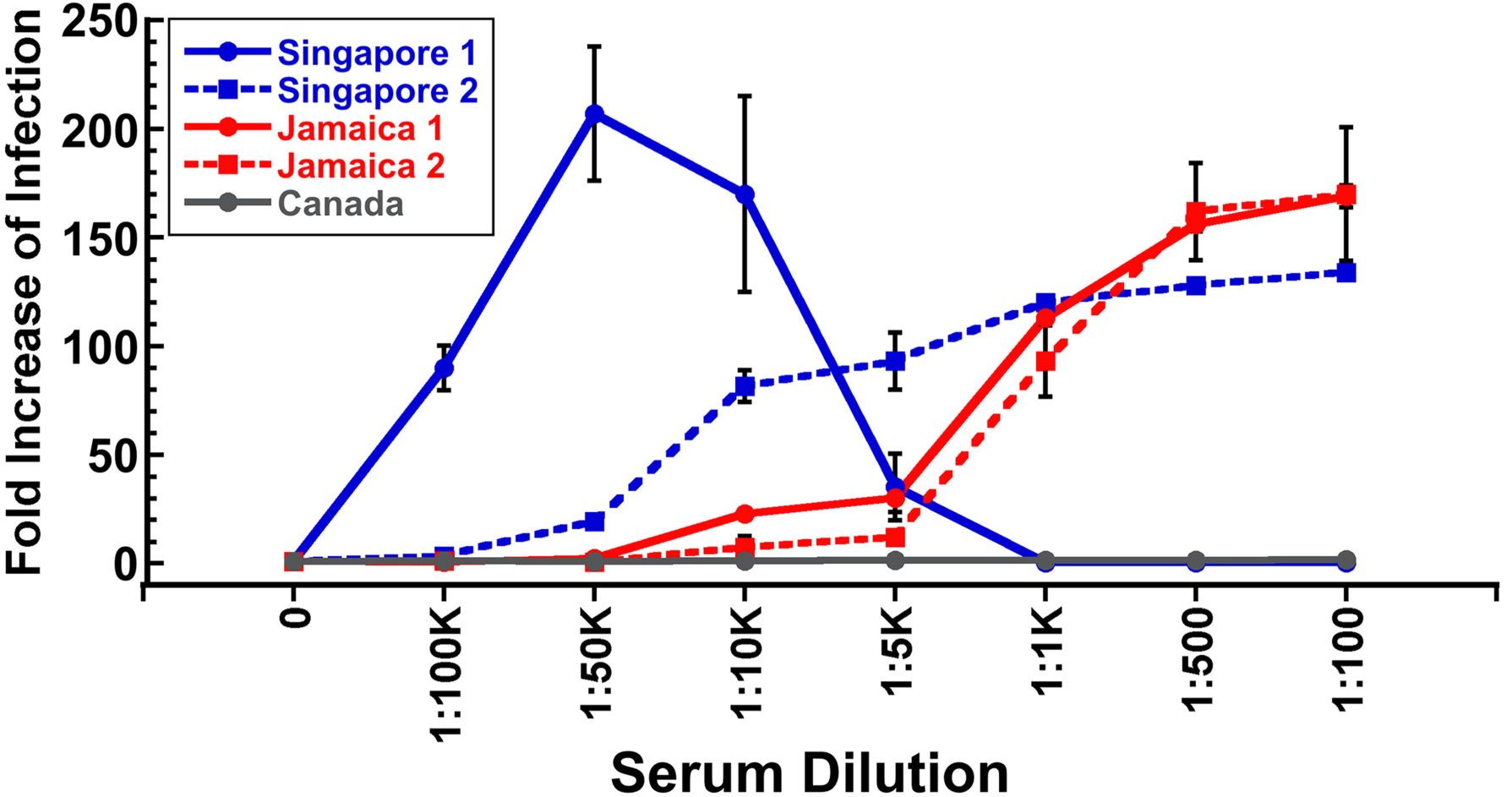
Enhancing activity of anti-DENV human sera against ZIKV. The effect of anti-DENV human sera on enhancement of ZIKV infection was determined in the human macrophage-like FcRII bearing cell line K562. All human anti-DENV sera tested showed strong infection enhancing activity of ZIKV MR766. At high serum concentrations, Singapore 1 serum blocked enhancement due to its strong neutralizing activity. Independent assays were repeated twice in triplicate.

**Fig 7.**
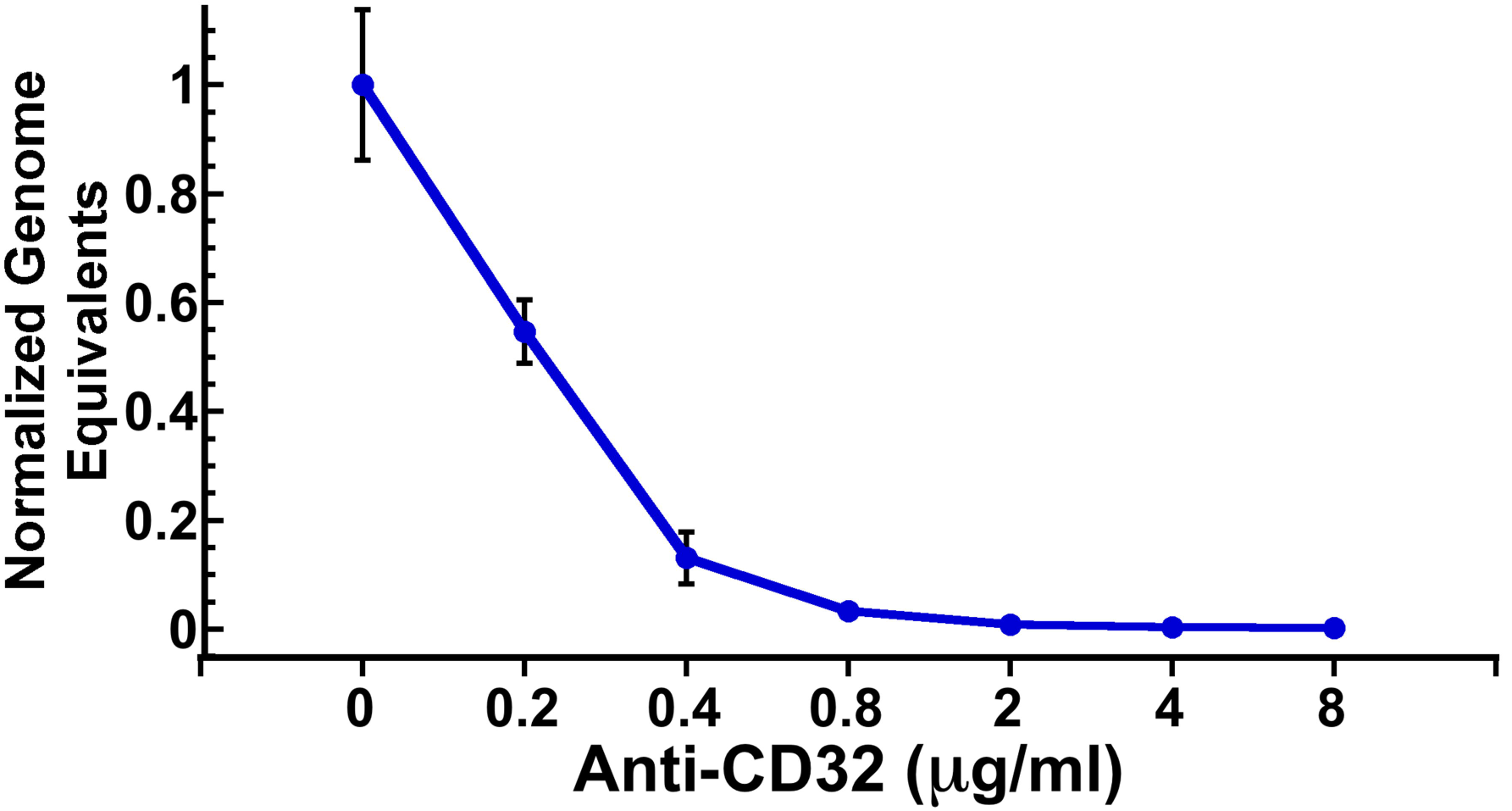
Anti-FcRII antibody blocks ZIKV enhancement activity of anti-DENV serum. K562 cells were pre-incubated with increasing concentrations of mouse anti-FcRII MAb prior to infection with ZIKV MR766 that had been pre-incubated with a highly enhancing dilution (1:50,000) of Singapore 1 serum. The results indicate that the ZIKV enhancement effect can be effectively blocked in a dose-responsive manner with an anti-FcRII MAb.

## Discussion

The present scenario of ZIKV introduction and spread in the Pacific and the Americas is complicated by pre-existing immunity to DENV. A recent serological survey of women giving birth in 2009-2010 in central Brazil documented that 53% of the new mothers were IgG positive for DENV [49]. ZIKV enhancement has been previously described to occur in the presence of cross-reactive sera raised against other flaviviruses. However, previous studies of ZIKV enhancement have not reported the effect of anti-DENV sera or antibodies or used human sera and cells [35,36]. Here we demonstrate that broadly neutralizing anti-DENV E protein fusion loop HMAbs cross-react with ZIKV, do not neutralize ZIKV, and greatly enhance ZIKV infection *in vitro*. Although the 10 amino acid E protein fusion loop region itself is identical between DENV and ZIKV, the binding epitope for these HMAbs is likely to be much larger and include important interactions with other variable portions of the E proteins that impact neutralization activity. We noted previously that these two HMAbs show little or no neutralizing activity against YFV or WNV [38].

In this study, we also investigated the role of secondary anti-DENV sera that might be considered as the worst-case scenario in DENV endemic regions. Our results show that human sera from secondary DENV infections can show varying degrees of neutralization, from neutralizing to non-neutralizing, and similarly enhance ZIKV infection. We have confirmed that the *in vitro* mechanism of ZIKV enhancement occurs through an FcRII-dependent process in human K562 cells in a manner very similar to DENV. If ZIKV ADE is fundamentally similar to DENV ADE, it is highly likely that preexisting anti-DENV antibodies will increase ZIKV viremia in humans and lead to more severe disease *in vivo*. This correlation will need to be confirmed clinically.

These results have implications for our understanding of ZIKV spread and persistence. In areas where DENV is endemic, ZIKV may transmit more readily and persist more strongly than expected from epidemiological transmission models of ZIKV alone, as has been observed in the recent ZIKV expansion in the Pacific and the Americas. How this plays out as ZIKV continues to spread in the Americas and other parts of the world where competent *Aedes* mosquito vectors are present, remains to be seen. One hopeful possibility is that ZIKV spread may be slower in areas where DENV immunity is low.

These results also have consequences for DENV and ZIKV vaccine design and use. We identified two serum samples that showed neutralizing activity against both DENV and ZIKV. The activity of highly neutralizing Singapore 1 serum is most likely explained by prior, undiagnosed ZIKV infection, whereas the Jamaica 1 serum neutralizing activity is likely not due to prior ZIKV infection, but may be a combined response against multiple DENV infections. In any case, this raises the possibility of inducing dual ZIKV and DENV immunity, perhaps with a single vaccine. Although the broadly neutralizing, anti-DENV HMAbs we tested did not neutralize ZIKV, there may be other human antibodies that may recognize and neutralize both ZIKV and DENV. However, DENV vaccines that induce a broadly reactive antibody response against viral surface envelope proteins with a large non-neutralizing antibody component may result in a cross-reactive, enhancing response against ZIKV, especially as the vaccine response wanes over time. Additionally, we know little about the reciprocal response of anti-ZIKV antibodies and their capacity to enhance DENV infections, although it would seem plausible that anti-ZIKV antibodies might similarly enhance DENV. A clear understanding of the interplay between ZIKV and DENV infections will be critical to ZIKV planning and response efforts in regions where ZIKV and DENV co-circulate, and particularly valuable for vaccine design and implementation strategies for both ZIKV and DENV.

## Acknowledgments

The authors would like to thank John S. Schieffelin at Tulane Univeristy for providing HMAbs 1.6D and D11C and Robert B. Tesh at the University of Texas at Galveston for providing virus strains through the World Reference Center for Emerging Viruses and Arboviruses.

